# Hox genes mediate the escalation of sexually antagonistic traits in water striders

**DOI:** 10.1101/425504

**Authors:** Antonin Jean Johan Crumière, Abderrahman Khila

## Abstract

Sexual conflict occurs when traits favoured in one sex impose fitness costs on the other sex. In the case of sexual conflict over mating rate, the sexes often undergo antagonistic coevolution and escalation of traits that enhance female’s resistance to mating and traits that increase male’s persistence. How this escalation in sexually antagonistic traits is established during ontogeny remains unclear. In the water strider *Rhagovelia antilleana*, male persistence traits consist of sex combs in the forelegs and multiple rows of spines and a thick femur in the rearlegs. Female resistance trait consists of a prominent spike-like projection of the pronotum. RNAi knockdown against the Hox gene *Sex Combs Reduced* resulted in the reduction of both the sex comb in males and the pronotum projection in females. RNAi against the Hox gene *Ultrabithorax* resulted in the complete loss or reduction of all persistence traits in male rearlegs. These results demonstrate that Hox genes can mediate sex-specific escalation of antagonistic traits along the body axis of both sexes.

## 1. Introduction

The evolutionary interests of males and females during mating interactions often diverge leading to the coevolution of sexually antagonistic traits that are favoured in one sex at a fitness cost to the other (Göran Arnqvist & Rowe, 2005; W. R. Rice & Holland, 1997). Empirical and theoretical data on the co-evolution of the sexes established sexual conflict as a major force in evolutionary change within and between lineages (G. Arnqvist, Edvardsson, Friberg, & Nilsson, 2000; Göran Arnqvist & Rowe, 2005). The consequences of sexually antagonistic selection are manifest in some spectacular shape changes in water striders, one of the most prominent model systems for the study of sexual antagonism in nature (G. Arnqvist & L. Rowe, 2002; Khila, Abouheif, & Rowe, 2012; Perry & Rowe, 2018). In many species, males are often favoured to mate repeatedly, but females pay increasing fitness costs for multiple mating (Göran Arnqvist & Rowe, 2005; Rowe, Arnqvist, Sih, & J Krupa, 1994). The repeated evolution of grasping traits that allow males to overcome female resistance is often matched by the evolution of anti-grasping traits in females that enhance their ability to resist (Göran Arnqvist & Rowe, 2005; Khila et al., 2012; G.A. Parker, 1979; G. A. Parker, 1983).

Sexually antagonistic traits are highly variable in shape and can occur in any segment along the body axis. Examples involve the modification of antennae, forelegs or rearlegs into grasping traits in the males, whereas females are known to match these with various anti-grasping traits such as erect abdominal spines (Goran Arnqvist & Locke Rowe, 2002; Khila et al., 2012; Westlake & Rowe, 1999). Several studies have highlighted the role of developmental genes in shaping certain male-specific morphologies in insects, including a case of an antagonistic trait in water striders (Khila et al., 2012; Ledon-Rettig, Zattara, & Moczek, 2017; Tanaka, Barmina, Sanders, Arbeitman, & Kopp, 2011; T. M. Williams et al., 2008). However, whether pleiotropic developmental genes can mediate the opposing escalation in antagonistic traits in both sexes have not yet been tested. Because Hox genes establish the identity of the segments along the body axis, we tested the role of two of these genes in the escalation of sexually antagonistic traits consisting of various sex-specific morphological modifications in a water strider species called *Rhagovelia antilleana* (Andersen, 1982; Crumiere et al., 2018; Moreira & Ribeiro, 2009; Polhemus, 1997). By inactivating *Sex Combs Reduced* (*Scr*) and *Ultrabithorax* (*Ubx*) during nymphal instars, we uncovered the importance of these two genes in shaping both male and female antagonistic traits.

## 2. Material and methods

### (a) Insect rearing

Laboratory populations of *Rhagovelia antilleana* were kept in water tanks at 25°C, 50% humidity, 14 hours of daylight and fed daily on crickets. Styrofoam floaters were provided for adult females to lay eggs on. Eggs were regularly transferred to separate tank to prevent cannibalism on the newly hatched nymphs.

### (b) Cloning of R. antilleana Scr and Ubx

Extraction of total RNA from *Rhagovelia antilleana* nymphs was performed using Trizol. cDNA was synthesized using SuperScript III First-Strand kit according to manufacturer’s instructions (Invitrogen). Fragments of *Scr* and *Ubx* genes were amplified by PCR using GoTaq G2 DNA polymerase (Promega) and the primers described in Table 1. PCR products were examined by electrophoresis, purified using PCR Minelute kit (Qiagen) and cloned into a pGEM-T vector kit (Promega). The sequences of Rhagovelia antilleana *Scr* and *Ubx* can be retrieved in GenBank using the following accession numbers: MG999826 and MG999808 respectively.

**Table 1:**
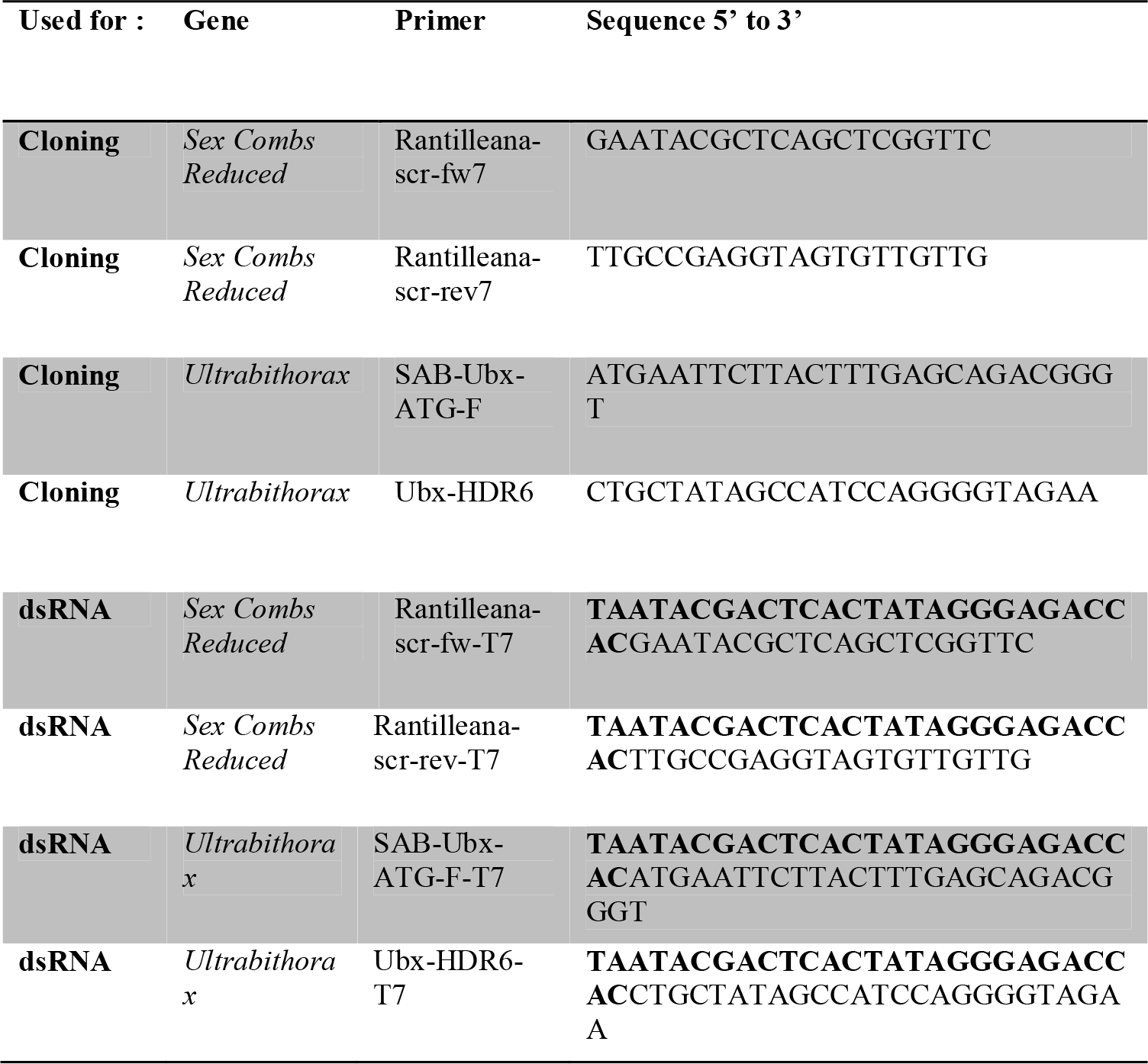
List of primers. Primers used to clone *Scr* and *Ubx* genes and to product dsRNA (T7 promoter sequence in bold) in *R. antilleana*.

### (c) Nymphal RNA interference in R. antilleana

A DNA template used to synthesize double-stranded RNA was produced using PCR on and *Scr* or *Ubx* primers tagged with a T7 RNA polymerase promoter and the *Scr* or *Ubx* plasmids as PCR templates (Table 1). The resulting PCR products were purified using Minelute kit (Qiagen) followed by an in vitro transcription with T7 RNA polymerase + (Ambion) to obtain the double stranded RNA corresponding to each gene (dsRNA). Both dsRNA were purified using RNeasy Mini Kit (Qiagen), concentrated with speedvac and re-suspended in 1X injection buffer (Rubin & Spradling, 1982) at a final concentration of 3 μg/μL. *Yfp* dsRNA was used as a control at 1,8 μg/μL concentration. We performed injections in *R. antilleana* first to third nymphal instars using a SteREO Discovery V8 (Zeiss), a Cell Tram Vario Oil Eppendorf injector and a Narishige micromanipulator under CO_2_ anaesthesia. The number of injected nymphs, emerged adults, and frequency of successful knockdown are presented in table 2.

### (d) Imaging

Image acquisition and observation of secondary sexual traits were performed using a SteREO Discovery V12 (Zeiss) and a ZEISS Merlin Compact Scanning Electron Microscope.

## 3. Results

### (a) *Scr* is required for both male persistence and female resistance traits

Secondary sexual traits in *R. antilleana* males and females only become prominent in the adult, thus indicating that these traits develop late during nymphal instars (Figure 1). Both males and females have evolved grasping and anti-grasping traits on the first thoracic segment. Male forelegs are equipped with a sex comb (Figure 2A) (Crumiere et al., 2018) whereas female pronotum exhibits a prominent spike-like projection (Figure 1 and Figure 2C) (Crumiere et al., 2018). Because the first thoracic segment in insects is under the control of the Hox gene *Scr* (Chesebro, Hrycaj, Mahfooz, & Popadic, 2009; Kopp, Duncan, Godt, & Carroll, 2000; Tanaka et al., 2011), we tested the role of this gene in the development of these structures. In males, *Scr* RNAi caused a notable reduction of the size of the teeth forming the comb (Figure 2B, Table 2). Interestingly in females, *Scr* RNAi also resulted in the reduction of the size and disruption of the shape of the pronotum projection (Figure 2D, Table 2). These results demonstrate that the same Hox gene, *Scr*, is involved both in the development of male persistence and female resistance traits that are located in its domain of action, *i. e*. the first thoracic segment.

**Figure 1:**
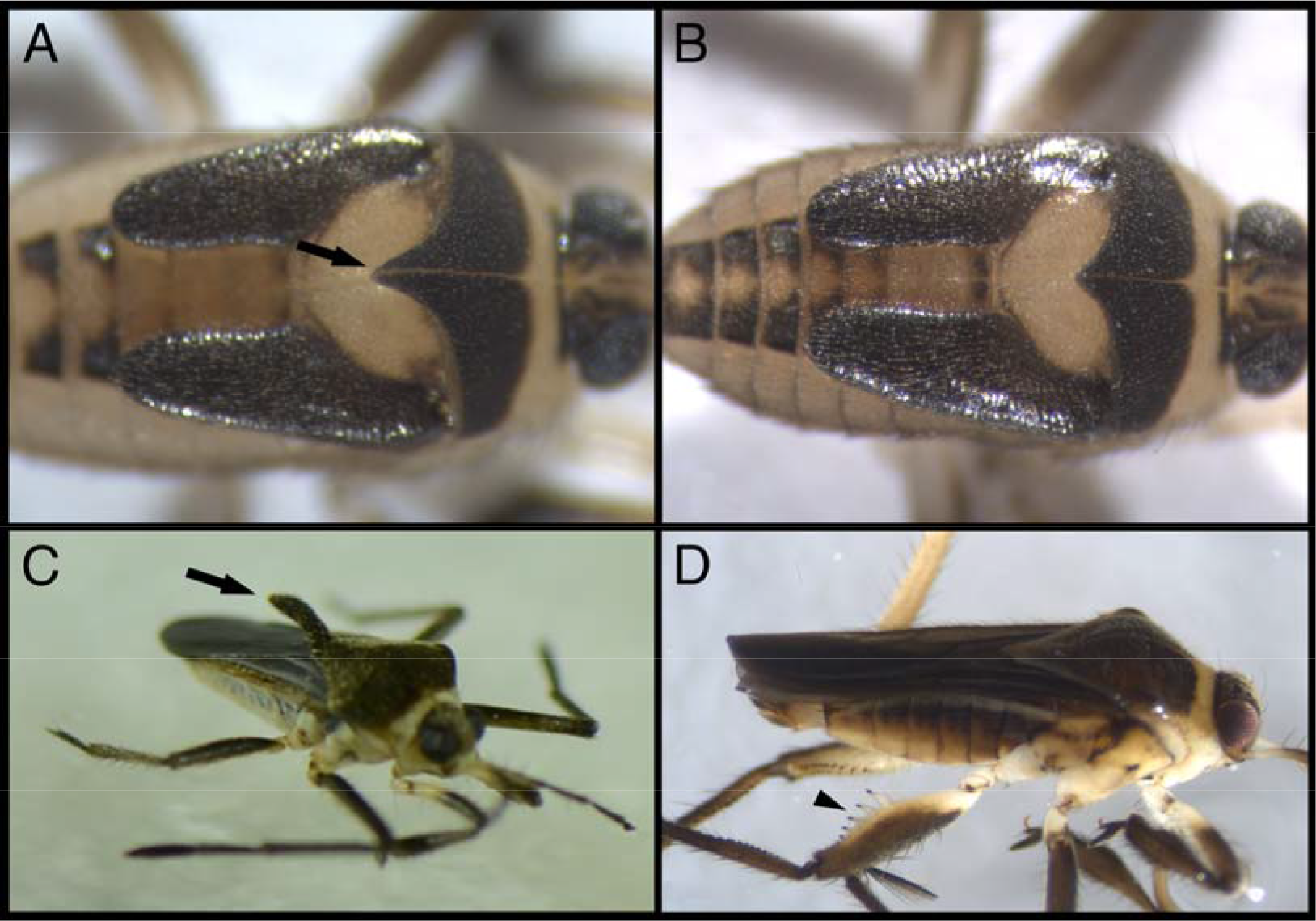
Development of some sexually antagonistic traits. **(A)** Fifth instar female nymph showing the extension of the pronotum as compared to male fifth instar nymph in **(B)**. **(C)** Adult winged female showing the pronotum projection, which is absent in adult males in **(D)**. Arrows point to the pronotum projection in females and arrowhead to male rearleg spikes.

**Figure 2:**
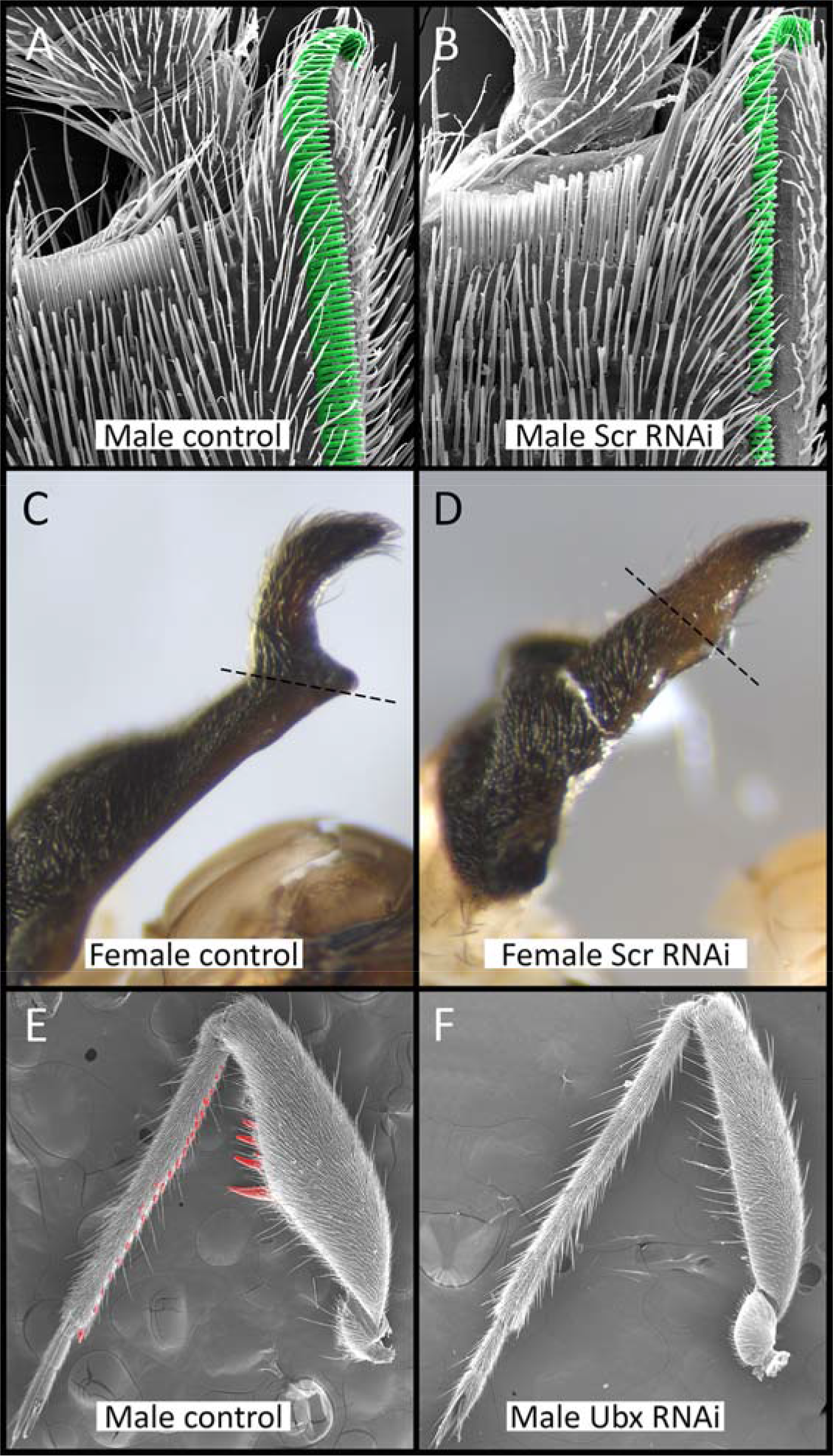
*Scr* and *Ubx* RNAi knockdowns phenotypes. In *R. antilleana* males, *Scr* RNAi induces a reduction of the size of the different teeth that composed the sex comb (A, B). In *R. antilleana* females, *Scr* RNAi induces the reduction of the size and a modification of the shape of the pronotum projection (C, D). In *R. antilleana* males, *Ubx* RNAi induces a reduction of the size of the rear-leg femur and a loss of the spikes present on the femur and on the tibia (E, F).

**Table 2:** Counting of nymph injections and phenotypes obtained in adults for *yfp* control, *Scr* and *Ubx* RNAi. Our RNAi experiment have a high level of penetrance for both *Scr* and *Ubx* with all adults obtained that show phenotypes compared to the control condition.

| Target gene | Number of injected individuals | Number of adult male obtained | Number of adult female obtained | Wild type phenotype obtained | RNAi phenotype obtained |
| --- | --- | --- | --- | --- | --- |
| *Yfp* (control) | 297 | 28 | 43 | 71 | 0 |
| *Scr* | 47 | 8 | 13 | 0 | 21 |
| *Ubx* | 348 | 22 | 27 | 0 | 49 |

### (b) *Ubx* shapes another set of male persistence traits found in the rearlegs

The rear-legs of *R. antilleana* males are equipped with sets of large and small spines arranged in rows on the trochanter, femur and tibia (Figure 1D and 2E) (Crumiere et al., 2018). We therefore tested the role of the Hox gene *Ultrabithorax*, which is known to specify the identity of the third thoracic segment in insects (Armisen et al., 2015; Davis, Srinivasan, Wittkopp, & Stern, 2007; Khila, Abouheif, & Rowe, 2014; Refki, Armisen, Crumiere, Viala, & Khila, 2014; Rozowski & Akam, 2002; Stern, 2003). Nymphal RNAi knockdown against *Ubx* resulted in the loss or reduction of all the armaments that otherwise develop on male rearlegs (Figure 1F, Table 2). Specifically, the width of the femur was significantly reduced and the spines on both femur and tibia were lost or reduced such that the rearleg of the male now resembles that of the female. This result indicates that *Ubx* mediates the development of male persistence traits located in the third thoracic segment in *R. antilleana*.

## 4. Discussion

We have shown that antagonistic coevolution of the sexes in *Rhagovelia* is developmentally controlled by the sex-specific action of the Hox genes *Scr* and *Ubx*. Interestingly the sex comb, which allows males to persist, and the pronotum projection, which allows females to resist (Crumiere et al. 2018), are both mediated by the same gene *Sex combs reduced. Ubx* on the other hand controls the elaboration of male rearlegs. How these genes can shape sex-specific morphologies during development remains unknown. Hox genes are known to control a large number of downstream targets that can diverge greatly among insects (Armisen et al., 2015; Pavlopoulos & Akam, 2011). It is possible that the sex-specific function of these Hox genes is mediated through sex-specific targets, as is the case for some fly sex-specific phenotypes (Barmina & Kopp, 2007; Thomas M. Williams & Carroll, 2009). The development of sexually antagonistic traits during late nymphal instars may have favoured changes in Hox targets to accumulate without any pleiotropic effects thus favouring rapid changes in Hox function. Other mechanisms could explain the sex-specific function of Hox genes in *Rhagovelia antilleana*. Interactions with sex-determination gene isoforms, such as *doublesex* (*dsx*) (G. Rice, Barmina, Hu, & Kopp, 2018; Tanaka et al., 2011; Wang & Yoder, 2012), would allow generating sex-specific structures by differentially regulating Hox genes (Barmina & Kopp, 2007; Tanaka et al., 2011), which in turn could also differentially regulate *dsx* in both sexes (Tanaka et al., 2011; Wang & Yoder, 2012). These interactions and regulations might allow Hox transcription factors being expressed in different cell populations within the same segment (Barmina & Kopp, 2007; Tanaka et al., 2011). Finally, following a similar mode of action of sex-specific isoforms for sex-determination genes (Ledon-Rettig et al., 2017) it would be interesting to investigate whether Hox genes express sex-specific isoforms and whether Hox sex-biased expression could be involved in antagonistic coevolution.

## Data accessibility

Additional data are available upon request.

## Author contributions

A.C. and A.K. conceived the study. A.C. performed experiment and collected and analysed the data. A.C. and A.K. interpreted the results and wrote the manuscript.

## Acknowledgments

We thank S. Viala for help with Scanning Electron Microscopy and the *Centre Technologiques des Microstructures* at Université Claude Bernard Lyon 1 for access to the Scanning Eletron microscope. This work was supported by an ERC Consolidator grant #616346 to AK.

## Ethical statements

All authors have agreed to be accountable for the accuracy and integrity of the work.

